# High throughput measurements of BMP/BMP receptors interactions using bio-layer interferometry

**DOI:** 10.1101/2020.10.20.348060

**Authors:** Valia Khodr, Paul Machillot, Elisa Migliorini, Jean-Baptiste Reiser, Catherine Picart

## Abstract

Bone morphogenetic proteins (BMP) are an important family of growth factors playing a role in a large number of physiological and pathological processes, including bone homeostasis, tissue regeneration and cancers. *In vivo*, BMPs bind successively to both BMP receptors (BMPR) of type I and type II, and a promiscuity has been reported. In this study, we used bio-layer interferometry to perform parallel real-time biosensing and to deduce the kinetic parameters (k_a_, k_d_) and the equilibrium constant (K_D_) for a large range of BMPs/BMPR combinations in similar experimental conditions. We selected four members of the BMP family (BMP-2, 4, 7, 9) known for their physiological relevance and studied their interactions with five type-I BMP receptors (ALK1, 2, 3, 5, 6) and three type-II BMP receptors (BMPR-II, ACTR-IIA, ACTR-IIB). We reveal that BMP-2 and BMP-4 behave differently, especially regarding their kinetic interactions and affinities with the type-II BMPR. We found that BMP-7 has a higher affinity for ACTR-IIA and a tenfold lower affinity with the type-I receptors. While BMP-9 has a high and similar affinity for all type-II receptors, it can interact with ALK5 and ALK2, in addition to ALK1. Interestingly, we also found that all BMPs can interact with ALK5. The interaction between BMPs and both type-I and type II receptors immobilized on the same surface did not reveal further cooperativity. Our work provides a synthetic view of the interactions of these BMPs with their receptors and paves the way for future studies on their cell-type and receptor specific signaling pathways.

## Introduction

Bone morphogenetic proteins (BMPs) are members of the transforming growth factor-β (TGFβ) superfamily who have been widely studied in view of the numerous physiological and pathological roles [1,2], including embryogenesis, development, bone homeostasis and regeneration and cancers [3]. The BMP family comprises more than 15 different ligands in humans, which have been grouped into four different subfamilies depending on their functions: BMP-2/4, BMP-5/6/7/8, BMP-9/10 and GDF5-6-7 [3–6].

Among these BMPs, BMP-2 is known for its role in morphogenesis, bone regeneration and musculoskeletal disorders [7,8]. In addition, BMP-4 plays a part in hematopoiesis and leukemia [9] while BMP-7 is involved in inflammation and glucose homeostasis [10]. BMP-9 and 10 have a major role in cardiovascular disease and anemia [11]. Furthermore, BMPs have been also reported to have an increasing role in cancer [12].

BMPs interact at the cell membrane with two sub-types of specific receptors (BMPR): type-I and type II BMPRs [2,4,5,13]. Seven different type-I receptors (ALK1 to ALK7) and four different type-II receptors (BMPR-II, ACTR-IIA, ACTR-IIB, TGFβR-II) are reported. BMPR-II, ACTR-IIA, ACTR-IIB associated to binding of all BMPs, while TGFβR-II are reported to be specific of TGFβ ligands, but not BMPs. BMPs have been reported to mostly bind to four receptors [3]: ALK1, ALK2 (also named ACTR-IA), ALK3 (also named BMPR-IA) and ALK6 (also named BMPR-IB). Each of these receptors has important physio-pathological roles. For instance, for the type-I BMPRs, ALK1 is the predominant receptor in endothelial cells and is involved in cardiovascular diseases [11]. ALK2 is an important receptor for bone homeostasis and a mutation in the ALK2 receptor is involved in a rare skeletal disorder named fibrodysplasia ossificans progressiva (FOP) and in a rare pediatric glioblastoma named diffuse intrinsic pontine glioblastoma (DIPG) [14], ALK3 plays a major role in several cancers, including breast and colorectal cancer [12], ALK6 plays a role in chronic myeloid leukemia [9,15]. ALK5 is reported to be a TGFβ receptor that is present in mesenchymal stem cells [16]. The three type-II receptors are usually considered to have similar roles in the signaling pathway associated to BMPs [3] but BMPR-II has likely been the most studied. Indeed, it was recently shown to play a protective role for endothelial cells from increased TGFβ responses and altered cell mechanics [17].

BMP signaling is initiated by the binding of BMPs to type-I BMPRs with high affinity prompting the constitutively active type-II BMPRs to come in close proximity to the formed complex, and induce the trans-phosphorylation of the glycine/serine-rich region (GS-box) preceding the kinase domain [1,18–20]. In this signaling pathway, the high number of BMP ligands (≈ 20) compared to the low number of BMP receptors (four type-I and three type-II receptors) indicates the presence of a promiscuous mechanism in which a given BMP can bind several receptors with distinct binding affinities [6][21].

Furthermore, it has been reported that high affinity ligands can compete with low affinity ligands for the binding of BMPRs and therefore can antagonize their signaling [22]. A better knowledge of the detailed binding characteristics of the BMPs to the BMPRs will help to identify the high affinity couples and to gain insight into the initiation of BMP signaling pathways.

To date, most of the characterizations of BMP/BMPR interactions have been determined using surface plasmon resonance (SPR) (23-37), which can be considered as a gold standard in the field. The data available from the literature of BMP/BMPR interactions are assembled in Table.1. However, the direct comparison of K_D_ for the different BMPs and BMP receptors is difficult since data has been obtained using various experimental conditions (different protein constructs, different immobilization strategies, different buffers, different SPR instruments…), which introduces a large variability in the experimental data. In addition, this data focused on particular BMP/BMPR couples and there is a lack of data for BMP-4 and 9 as well as for ALK1.

**TABLE 1.**
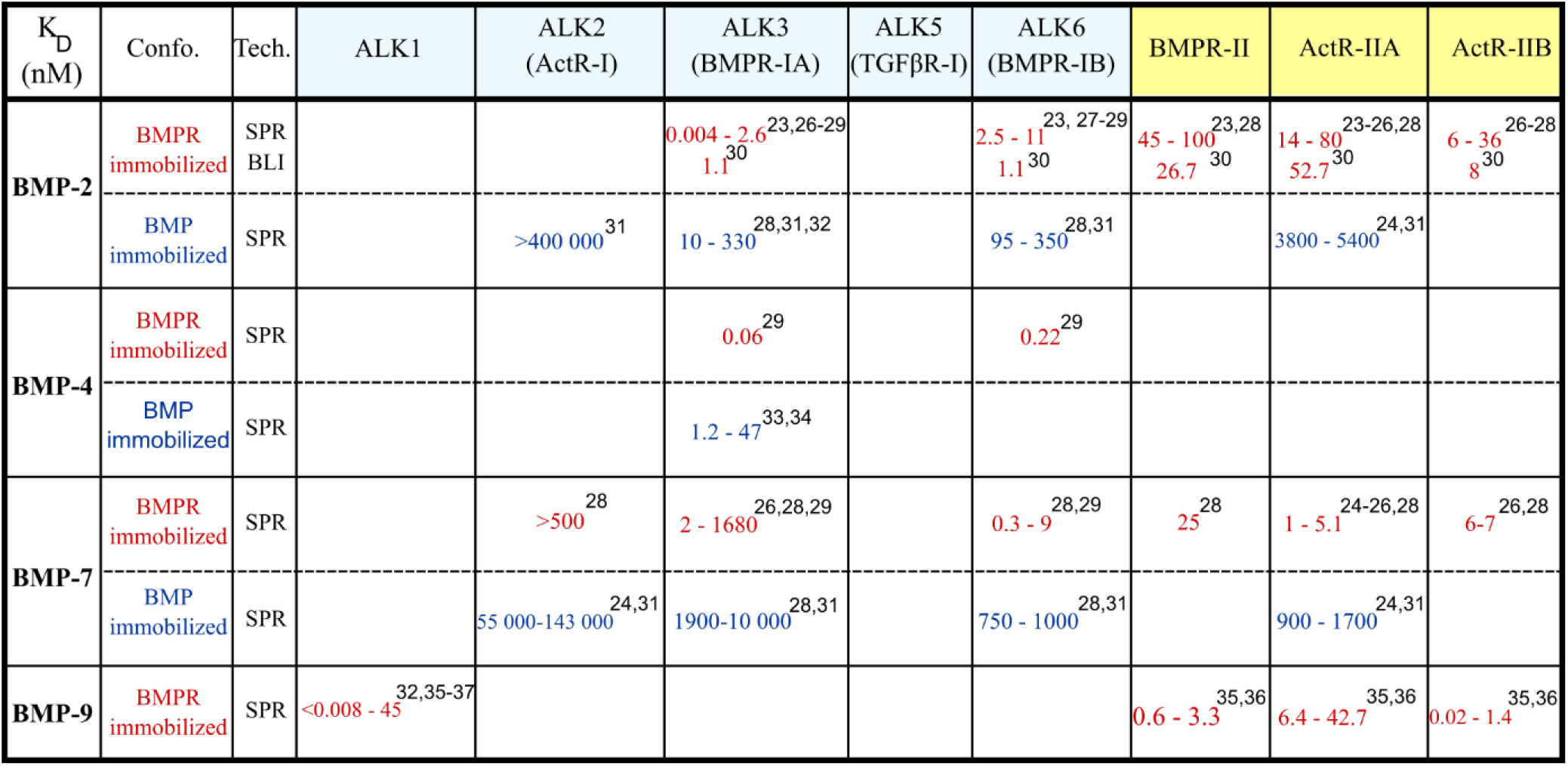
Literature study of all the K_D_ (nM) of the interactions between couples of BMP and their receptors (BMPR of type I and type II) The experiments were usually performed in one of the two configurations: in red, when the BMPR is immobilized and in blue, when the BMP is immobilized.

Among all the biophysical methods available today to characterize protein-protein interactions, the reflectometric interference spectroscopy (RifS) [38,39] is a very recent label-free optical method based on white light interferences at layers of sensors. A commercially-available setup known as bio-layer interferometry (BLI) enables to perform parallel real-time binding measurements and characterization of biomolecule interactions. It is increasingly used to study kinetic constants and binding affinities of protein-protein and protein-nucleic acid interactions [39–42] and it has only recently begun to be used to study BMP/activin A chimera interactions [30].

In the present study, our aim was to quantify in similar experimental conditions and a large set of BMP/BMPR interactions in a parallel manner, in order to directly compare their kinetic parameters and binding affinities. We decided to focus on four BMPs that are among the most widely studied: BMP-2, BMP-4, BMP-7 and BMP-9 [43]. For the type-I BMP receptors, we considered ALK1, ALK2, ALK3, ALK6 and added ALK5 known as an essential TGFβ receptor [44], since it is involved in the signaling of BMP-responsive cells such as mesenchymal stem cells [16]. We studied the three type-II BMP receptors (BMPR-II, ACTR-IIA and ACTR-IIB).

## Results

We first performed a literature study to gain information on the state-of-the-art regarding BMP/BMPR interactions. Table.1 provides a view of the K_D_ values, which are in the nM range for the highest affinity interactions. Table.SI.1 gives the detailed information obtained from each published study. We first note that all experiments, but one using the commercially-available BLI setup [30], were conducted using SPR with two configurations to perform the experiments: the first configuration consists in immobilizing the BMPs on the sensor chip while the second consists in immobilizing the BMPRs, this second strategy being the most common. In terms of immobilization protocols, we noted that several strategies were proposed, which can be grouped in three major categories (Table.SI.1): i) using biotinylated BMPR coupled to streptavidin-coated surfaces; ii) using BMPR-Fc captured on anti-Fc coated sensors and iii) direct immobilization of BMPR using an amine coupling strategy.

Looking at the published studies (Table.1 and Table SI.1), it appears from that for a given BMP/BMPR couple, the range of measured K_D_ can be very broad. These discrepancies likely arise from the differences in experimental details, including immobilization strategies, experimental working conditions and the biochemistry of BMPs itself. Moreover, since BMP-2 and BMP-4 are usually considered to behave similarly [3], several studies were performed only on BMP-2 interaction with type-II BMPRs (BMPR-II, ACTR-IIA and ACTR-IIB) and with type-I BMPR ALK2, but there is no such study for BMP-4. We also noted a lack of data for the interactions of BMP-2, 4 and 7 with ALK1 since it was reported to be the major BMP-9 receptor [11]. Lastly, we noticed the absence of data on ALK5 (TGFβR-I) with any of the chosen BMPs, since it was traditionally considered solely as a TGFβ receptor [44] but was also shown to be a central point in BMP/TGFβ signaling [45].

### Dimeric state of BMPs and BMPR

The commercially-available proteins that we used were produced in CHO for BMP-2, 7 and 9 or in *E.coli* for BMP-4. The BMPR coupled to Fc fragments (BMPR-Fc) were produced in mouse myeloma NS0 cells, except for ACTR-IIA that was produced in CHO cells. We verified the biochemical state (monomeric or dimeric) of all BMPs and BMPR-Fc by gel electrophoresis in both non-reducing and reducing conditions (Fig.SI.1). The BMPs were mostly dimeric, as expected [6], and migrate at ≈ 26 kDa in non-reducing conditions, and at ≈ 13 kDa in reducing conditions. Since the Fc fragment form dimers, the BMPR chimeras are also present in dimeric state and migrate at 90 and 110 kDa in non-reducing condition and in a monomeric form with a band between 45 and 55 kDa in reducing conditions (Fig.SI.1).

### Immobilization of BMPR on the biosensor

*In vivo*, the BMPs are soluble proteins that localize in the extracellular matrix or in blood for BMP-9. They can then be considered to diffuse freely in a 3D space. The BMPRs are trans-membrane proteins that are localized at the cellular membrane and are thus diffusing in a 2D space. For this reason, it is likely that the order of magnitude of the diffusion of BMPR is similar to that of lipids in a membrane (≈ 1 µm^2^/s) while that of BMPs is similar to a protein diffusing freely (≈ 100 µm^2^/s) [46]. We thus choose to immobilize the BMPRs at the biosensor surface and to adsorb BMPs at their surface to better mimic the *in vivo* situation.

In order to find a protocol applicable to all BMP/BMPR couples, we considered several capture strategies for BMPR immobilization at the biosensor surface. The same previously-published capture methods used for SPR, including biotinylated ligand/streptavidin surface, amine coupling absorption or Fc chimera/anti-human IgG or protein A surfaces were considered (Table.SI.1) [23–25,27]. Since all the BMPR-Fc chimeras were commercially-available, and since anti-Fc fragment-coated biosensors are known to more stable than protein A [30,35], we selected this strategy that consists in immobilizing the BMPR-Fc chimeras via the Fc dimer to the biosensor surfaces (Fig 1A and Material and methods). This configuration presents the advantage of immobilizing all of the BMPR homogeneously in one orientation, with their binding site accessible to BMPs.

**FIGURE 1.**
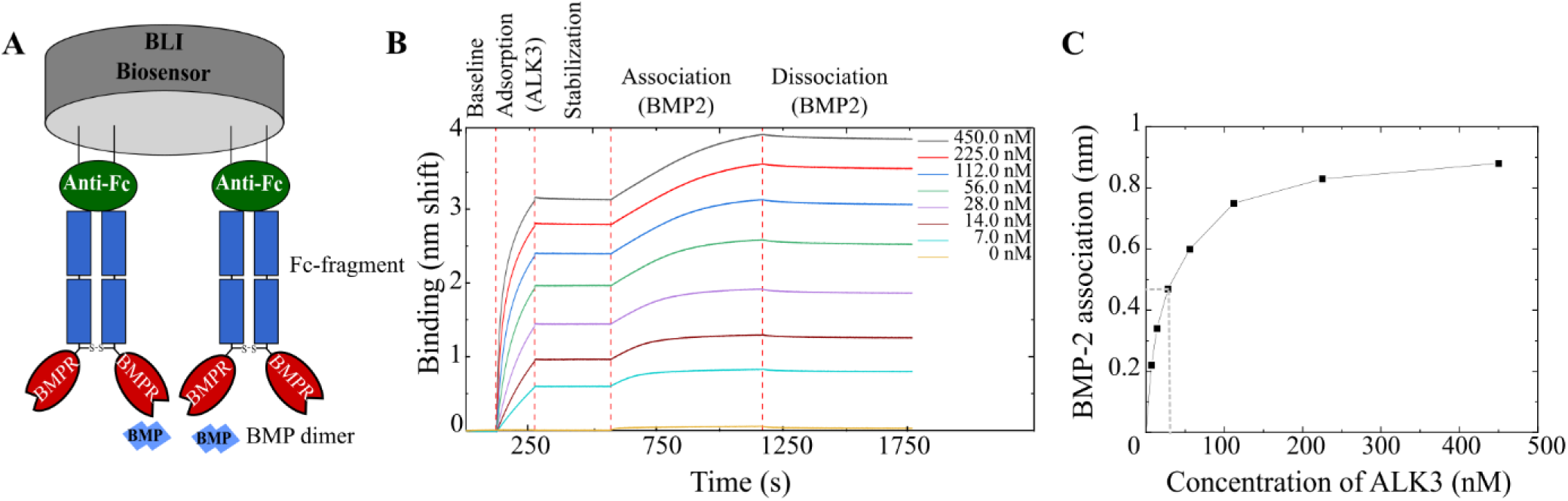
Adsorption strategy of BMPR-Fc on the biosensors. **A)** Schematic representation of the adsorption strategy where an anti-Fc-coated biosensor binds the Fc-Receptor chimera. **B)** Preliminary experiment where ALK3 receptor was adsorbed at increasing concentrations (from 7 nM to 450 nM) and set to interact with BMP-2 at a concentration of 5 nM. **C)** The interaction signal of BMP-2 to ALK3 given in nm shift, plotted as a function of ALK3 initial concentration in solution.

In order to determine a suitable adsorption density of the BMPRs on the biosensors, we performed preliminary assays with ALK3 receptor immobilized at increasing densities leading to a signal between 0.5 to 3 nm of spectral shift (nm) after a fixed contact time of 150 s. The functionalized surfaces were then set in contact with BMP-2 at a constant concentration of 5 nM to proceed to BMP-2 adsorption (Fig.1B). As shown by the response at equilibrium *vs.* ALK3 concentration (FIG.1C), the optimal concentration of ALK3 is ≈ 28 nM, leading to an association signal of ≈ 0.5 nm after 600s.

### Interaction of BMPs with the BMPR-I and BMPR-II

The kinetic interaction studies were then performed using the same protocol for the four BMPs with BMPRs. All BMPRs were adsorbed at densities corresponding to a spectral shift between 0.8 and 1.1 nm. The BMP concentrations were varied over a large range ranging from 2 nM to 80 nM (Fig.2). Representative experimental curves for BMP-2/ALK3, BMP-9/ALK1, BMP-2/BMPR-II and BMP-7/ACTR-IIA are shown in Fig.2 (respectively panel A-D).

**FIGURE 2.**
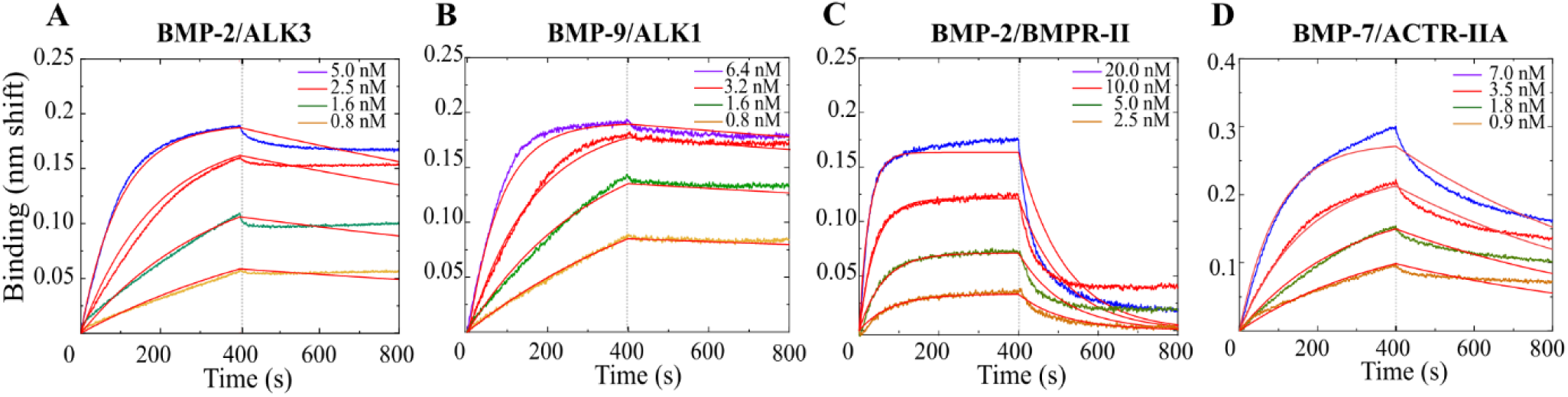
Examples of binding kinetics between type-I BMPRs and BMPs. **A)** BMP-2/ALK3, **B)** BMP-9/ALK1 and between type-II BMP and BMPs, with **C)** BMP-2/BMPR-II and **D)** BMP-7/ACTR-IIA.

To determine the kinetic parameters, the 1:1 Langmuir model binding model has been used. Indeed, it has been shown in structural studies [23,25,47] that the BMP/BMP receptor interaction can be considered as bimolecular: It was reported that BMP dimers comprise two distinct pairs of binding sites: one for type-I BMPR and the other for type-II BMPR. While the type I interface is a large continuous area formed by residues from both BMP monomers, the interface with type II is composed only of amino-acids from one BMP monomer [5], as seen in the example of BMP-2/ALK3/ACTR-IIA (pdb: 2H64) [48] (Fig.3.A). Thus, a one to one binding is expected. Furthermore, the commonly-used 1:1 Langmuir model has always been used to date to fit the experimental data and to determine kinetics parameters [23,26,49]. In the present case, since the Fc chimera induces a dimerization of the BMPR, two possible binding modes are possible (Fig.3.B-E): one BMP molecule binding to two proximate BMPR binding domains (model B or D) or one BMP molecule binding to one BMPR binding domain (model C or E). Nonetheless, since BMP dimers are fully symmetrical, all binding models may lead to 1:1 binding kinetic. The R2 values of the fits, presented in tables SI.2 and SI.3, are in majority around 0.95 and higher which indicates an acceptable fit.

**FIGURE 3.**
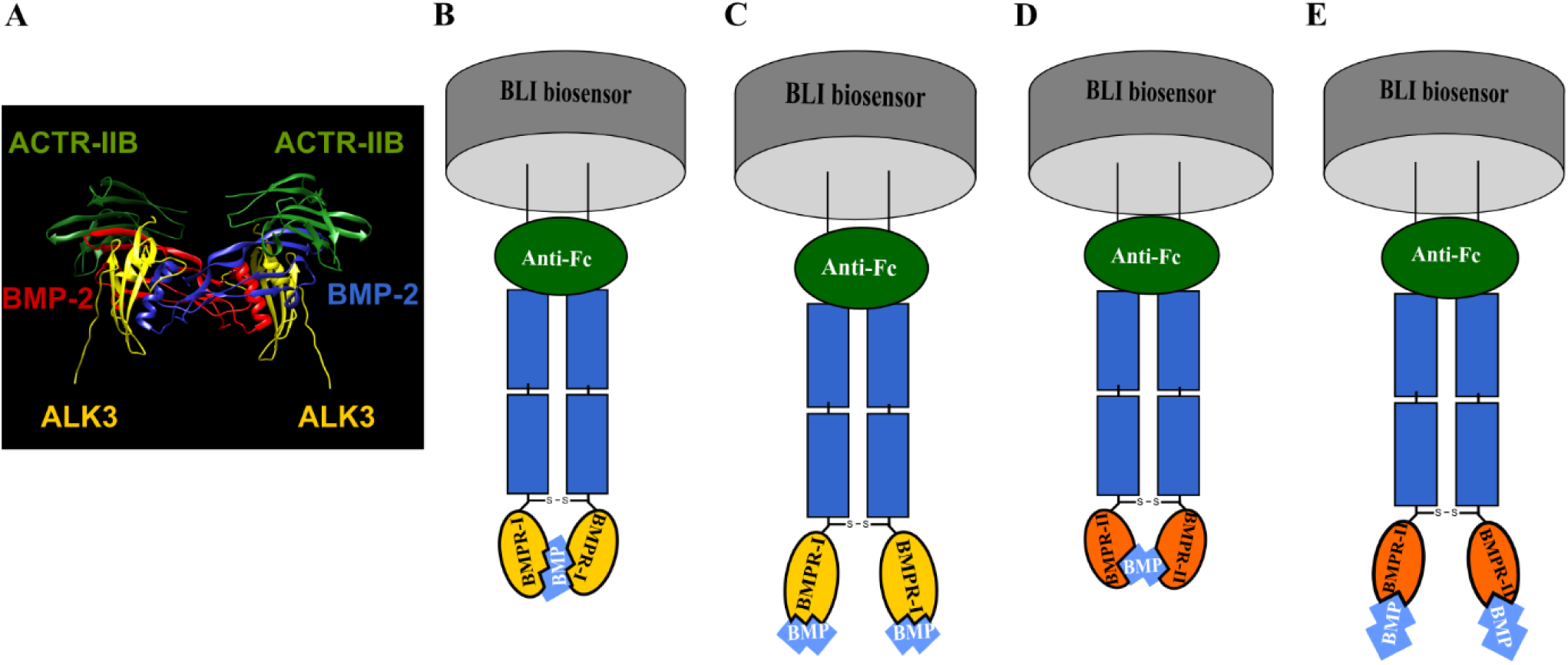
A schematic representation of the different binding models in simple BMP/BMPR interaction. **A)** Picture of BMP-2/ALK3/ACTR-IIB (pdb:2H64) adapted from Weber, D. et al ***BMC struct Biol*** 2007 [48]. **B)** Association of the two binding sites of BMP dimer to two type-I BMPR. **C)** Association of one binding site of BMP dimer with one type-I BMPR. **D)** Association of the two binding sites of BMP with two type-II BMPR. **E)** Association of one binding site of BMP dimer with type-II BMPR.

**FIGURE 4.**
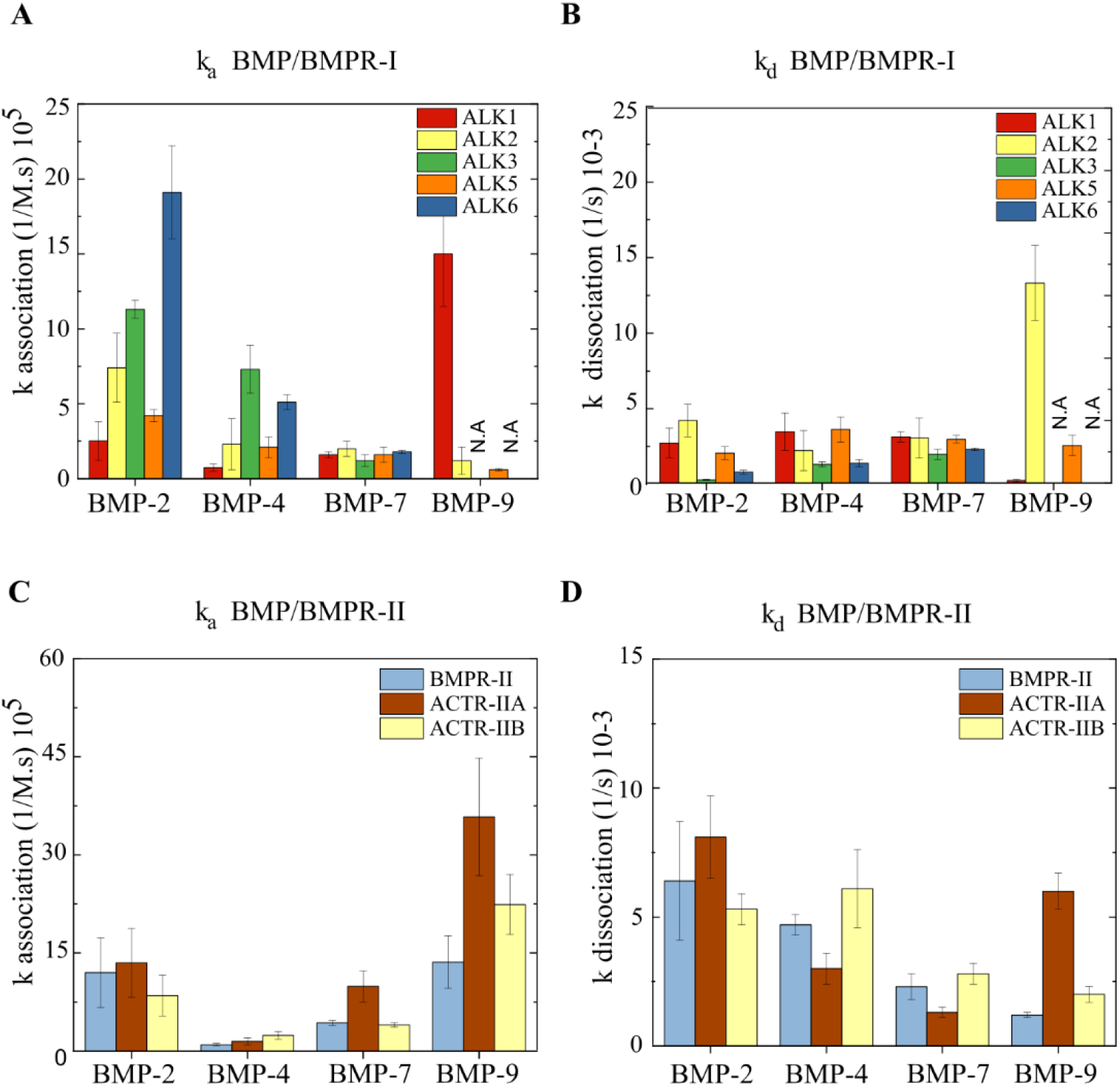
Histograms presenting the association constants (k_a_) and the dissociation constants (k_d_). k_a_ of **A)** BMP/type-I BMPR and **B)** BMP/type-II BMPR interactions and k_d_ of **C)** BMP/type-I BMPR and **D)** BMP/type-II BMPR interactions. For BMP-9/ALK3 AND BMP-9/ALK6, the signal was very low (N.A). The error bars represent the s.d (n=3).

A fast association was generally observed for all the BMPs interacting with the type-I BMPR, but differences in the dissociation rate are seen. The association constant (k_a_) and dissociation constant (k_d_) that were extracted from the fit of each interaction curve are presented in Fig.4 as well as tables SI.2 and SI.3. BMP-2 and BMP-4 exhibit a high k_a_ (≈ 15×10^5^ M^−1^.s^−1^ for BMP-2 and ≈ 5×10^5^ M^−1^.s^−1^ for BMP-4), and low k_d_ with both ALK3 and ALK6 (≈ 0.5×10^−3^ s^−1^ for BMP-2 and ≈ 1.5×10^−3^ s^−1^ for BMP-4), indicating a fast association and slow dissociation to these receptors. Furthermore, BMP-2 associates and dissociates in a similar manner to ALK1, ALK2, ALK5 (k_a_ ≈ 4×10^5^ M^−1^.s^−1^ and k_d_ ≈ 3×10^−3^ s^−1^) and to the three type-II BMPRs (k_a_ ≈ 11×10^5^ M^−1^.s^−1^ and k_d_ ≈ 6×10^−3^ s^−1^). In comparison to BMP-2, BMP-4 associates more slowly to these receptors.

BMP-7 demonstrates a slow association to all the type-I BMPRs (≈ 2 x10^5^ M^−1^.s^−1^), in addition to a slow association (≈ 6×10^5^ M^−1^.s^−1^), and a slow dissociation (≈ 2 x10^−3^ s^−1^) to type-II BMPRs. Regarding BMP-9, it exhibits a fast association (15.0 ± 3.5 x10^5^ M^−1^.s^−1^) and a very slow dissociation (0.2 ± 0.1 x10^−3^ s^−1^) to ALK1 and type-II BMPR (k_a_ ≈ 20 x10^5^ M^−1^.s^−1^ and k_d_ ≈ 3.3 ± x10^−3^ s^−1^). BMP-9 also presents a slow association and fast dissociation from ALK2 and ALK5 (k_a_ ≈ 2×105 M^−1^.s^−1^ and k_d_ ≈ 3×10^−3^ s^−1^), but it does not interact with ALK3 and ALK6.

Next, we calculated the equilibrium affinity constant K_D_ (equal to the ratio of k_d_ over k_a_). BMPs present a generally high affinity to all BMPRs ranging from 133 to 0.2 nM for high affinity interactions. The lowest K_D_ values are highlighted in dark blue. BMP-2 and BMP-4 have a good binding affinity to both ALK3 and ALK6 since their K_D_ was < 3 nM (Table.2A). They bind to ALK2 similarly with an affinity of 7.0 ± 2.3 nM for BMP-2 and 10.5 ± 3.8 nM for BMP-4. They also bind to ALK1 and ALK5 but BMP-2 has a ≈ 4-fold higher affinity to these receptors than BMP-4. Regarding type-II BMPRs, BMP-2 had a similar affinity for both ACTR-IIA and ACTR-IIB (≈ 6 nM) while BMP-4 also interacted with both receptors although with ≈ 4-fold lower affinity (≈ 23 nM). In addition, BMP-2 has also a 10-fold higher affinity for BMPR-II than BMP-4. We then investigated whether the differences between BMP-2 and BMP-4 may arise from their glycosylation state, since BMP-2 is produced in CHO while BMP-4, being produced in *E-Coli*, is non-glycosylated. We thus compared the interactions of ALK3 with both the glycosylated and non-glycosylated forms of BMP-4 (Fig.SI.2). For the glycosylated form of BMP-4, the increase in the non-specific signal was negligible (≈ 0.02 nm). However, the interactions differed slightly since K_D_ was 1.32 ± 0.48 nM for the non-glycosylated BMP-4, versus 0.3 ± 0.06 nM for the glycosylated form.

**TABLE 2.** Binding affinities (K_D_ in nM) of BMP/BMPR interactions. Tables summarizing the K_D_ (nM) of the BMP/BMPR interactions for: **A)** type I and **B)** type II receptors, obtained from the kinetic experiments in a conformation where the BMPR is immobilized. The interactions between BMP-9/ALK3 and BMP-9/ALK6 yielded a very low signal (N.A). The high affinity couples are colored in dark blue. The error values represent the s.d (n=3).

**A**
| BMPR<br>BMP | ALK1 | ALK2<br>(ActR-I) | ALK3<br>(BMPR-IA) | ALK5<br>(TGFβR-I) | ALK6<br>(BMPR-IB) |
| --- | --- | --- | --- | --- | --- |
| BMP-2 | 13.0 ± 1.6 | 7.0 ± 2.3 | 0.21 ± 0.03 | 5.8 ± 1.1 | 0.5 ± 0.1 |
| BMP-4 | 55.4 ± 4.0 | 10.5 ± 3.8 | 1.7 ± 0.5 | 21.9 ± 6.6 | 3.1 ± 0.3 |
| BMP-7 | 23.1 ± 2.1 | 18.4 ± 3.8 | 19.0 ± 2.1 | 22.6 ± 1.1 | 15.0 ± 1.2 |
| BMP-9 | 0.2 ± 0.1 | 133.1 ± 35.1 | N.A | 51.0 ± 18.3 | N.A |

**B**
| BMPR<br>BMP | BMPR-II | ACTR-IIA | ACTR-IIB |
| --- | --- | --- | --- |
| BMP-2 | 5.4 ± 0.8 | 6.1 ± 1.2 | 6.3 ± 3.4 |
| BMP-4 | 56.0 ± 6.0 | 21.4 ± 3.7 | 26.0 ± 0.5 |
| BMP-7 | 5.5 ± 1.2 | 1.3 ± 0.3 | 7.1 ± 0.7 |
| BMP-9 | 0.8 ± 0.2 | 1.7 ± 0.1 | 1.4 ± 0.4 |

BMP-7 interacts with all type-I BMPRs with similar affinities (≈ 20 nM). In contrast, it has a greater affinity for the three type-II BMPRs as it binds to BMPR-II and ACTR-IIB similarly (≈ 6 nM) and to ACTR-IIA with a 5 to 7-fold higher affinity (1.3 ± 0.3 nM).

Regarding BMP-9, it binds ALK1 with high affinity (0.2 ± 0.1 nM), ALK5 and ALK2 with a much lower affinity (51.0 ± 18.3 nM and 133.1 ± 35.1 nM, respectively). The affinity of BMP-9 for all the three type-II BMPRs is high: 0.8 ± 0.2 nM for BMPR-II, 1.7 ± 0.1 nM for ACTR-IIA and 1.4 ± 0.4 nM for ACTR-IIB. Notably, BMP-9 affinity for BMPR-II is about 2-fold higher in comparison to ACTR-IIA and ACTR-IIB.

Thus, the K_D_ values indicated that there are significant differences between BMP-2 and BMP-4, a higher affinity of BMP-7 to type-I BMPRs in comparison to type-I BMPR, and a highly selective affinity of BMP-9 for ALK1 as well as to all type-II BMPRs.

In order to compare the BLI technique to SPR (Table.1), we performed SPR kinetic experiments for selected high affinity couples, namely BMP-9/ALK1 and BMP-2 or BMP-4/ALK3 or ALK2. For this purpose, we used commercially-available protein A-coated chips and BMPR-Fc chimera as adsorption strategy (Fig.5A). Unfortunately, the BMP-2/ALK3 (Fig.SI.3), BMP-2/ALK2, BMP-4/ALK3 (data not shown) kinetic interaction using this adsorption strategy could not be measured since non-specific binding to the sensor ship was too high and specific binding signal could not be resolved (Fig.SI.3). In contrast, the BMP-9/ALK1 interaction was notable and a K_D_ of 13.4 pM was obtained. This value is 15-fold lower than obtained by BLI (~200 nM – Fig.5B).

**FIGURE 5.**
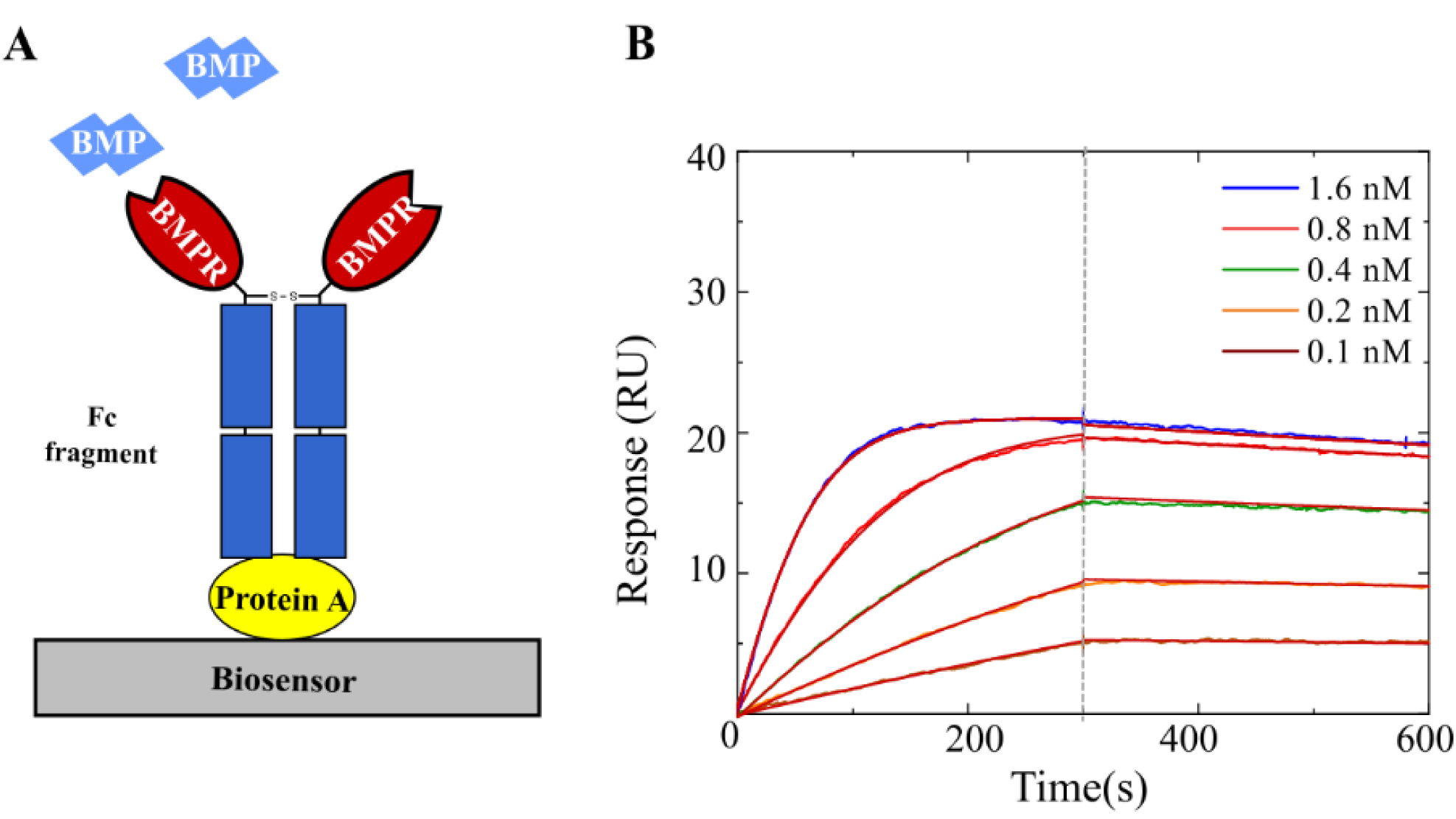
SPR study of BMPR/BMP interactions. **A)** A schematic representation of a SPR biosensor surface where the interaction was studied. Protein A was used to immobilize the BMPR Fc fragment. **B)** Example of kinetic experiment for the ALK1/BMP-9 couple showing the association phase (up to 300s followed by the dissociation phase).

### Interaction of BMPs with the BMPR-I/BMPR-II ternary complex

Next, we decided to investigate the interactions of BMP to type-I/type-II BMPR complexes. *In vivo*, It is reported that BMPs first bind to the inactive type-I BMPRs thus triggering type-II BMPRs to activate (by phosphorylation) the type-I BMPRs by forming a ternary complex [24,50]. We studied ALK2 as a type-I BMPR and all three type-II BMPRs with BMP-2, BMP-4 and BMP-7. We chose ALK2 since it is the most studied receptor and has a middle range affinity for the BMPs. Our experimental approach consisted in loading sequentially both types of BMP receptors on the biosensor (Fig.6A-B). We performed experiments using two capture strategies. Firstly, ALK2 was loaded, followed by a type-II BMPR and then BMP-2 was set into contact with the functionalized surfaces (Fig.6A). Secondly, a reverse sequence was used in which ACTR-IIB was captured first, followed by ALK2 and then BMP-2 (Fig.6B). The adsorption times were chosen such as to have an equivalent level of adsorption for each receptor, with a total shift being similar to the case of single BMP/BMPR interactions.

**FIGURE 6.**
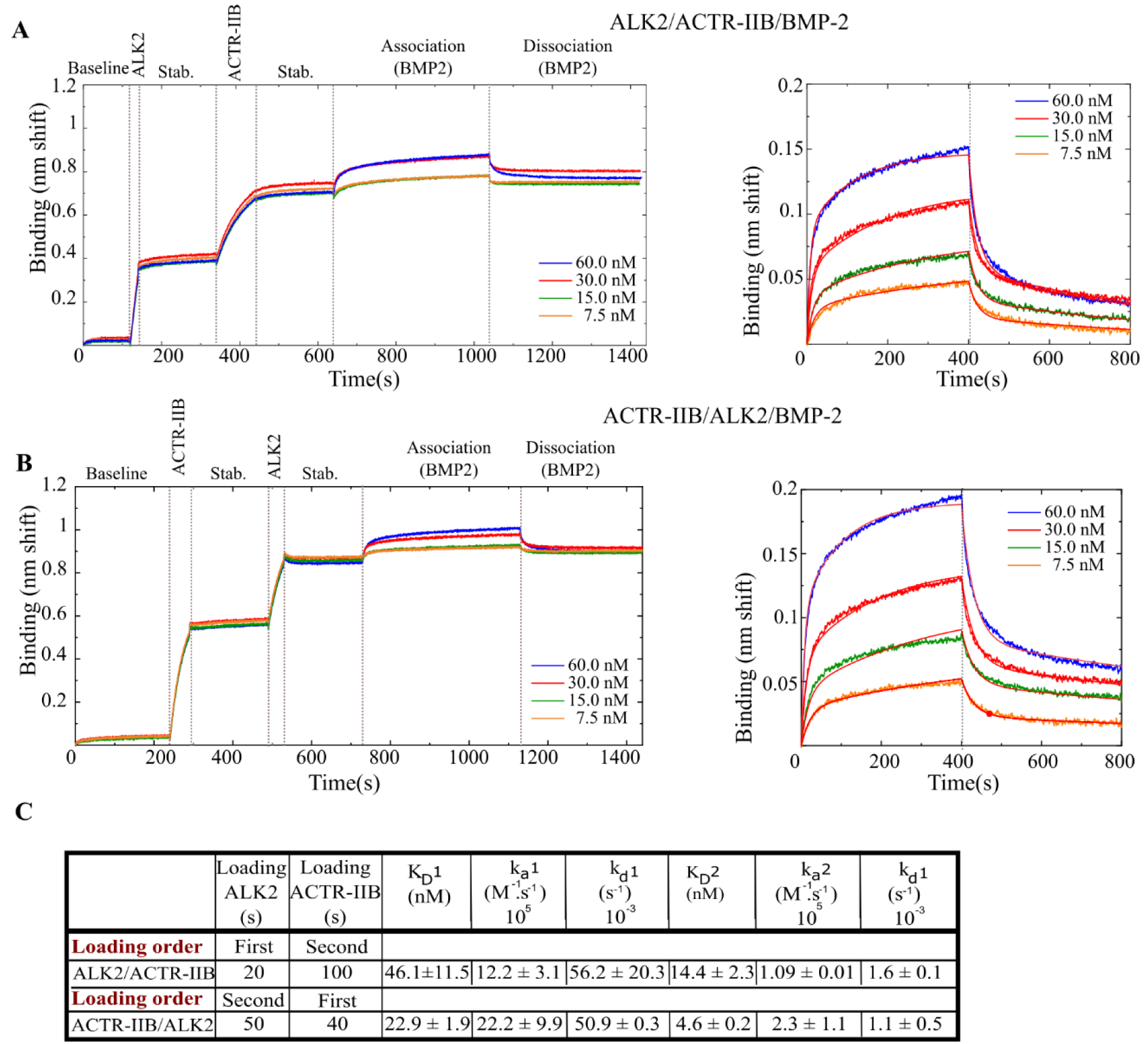
Interaction between the ALK2/ACTR-IIB heterocomplex and BMP-2. Binding was done sequentially: ALK2 or ACTR-IIB first followed by the second receptor and then BMP-2. **A)** First ALK2 or **B)** first ACTR-IIB. **C)** A representation of a ternary complex model. **D)** Table summarizing the kinetic parameters deduced from the experimental fit to the data. The stab. step refers to stabilization.

To process and fit the kinetic data, we initially applied a 1:1 Langmuir model but the fit was of poor quality (Fig.SI.4). These interactions consist of two different receptors and therefore possess two pairs of two distinct binding sites for type-I and type-II BMPRs. We thus presumed that both types of type-I and type-II receptors bind BMPs separately with different affinities and therefore applied a 2:1 heterogeneous binding model.

The K_D_ in the configuration where ALK2 was immobilized first was 46.1 ± 11.5 nM for the first binding site and 14.4 ± 2.3 for the second. Conversely, when BMPR-II was adsorbed first the K_D_1 was 22.9 ± 1.9 nM and K_D_2 4.6 ± 0.2 nM (Fig.6D). The same experiment was also performed for ALK2/BMPR-II/BMP-7 resulting in a K_D_ of 4.6 ± 0.7 nM and 19.1 ± 6.4 nM for the experiment where ALK2 was adsorbed first, compared to 5.4 ± 0.2 nM and 23.2 ± 0.1 nM for the reverse order. In the case of BMP-7, binding is similar whatever the order of receptor presentation.

The binding affinities for all the experiments where ALK2 is loaded first are summarized in Table.3. Surprisingly, we did not find any improvements in the K_D_ when two receptors are captured on the biosensor surfaces compared to the situation when only one is present. The K_D_ for all of the experiments appear to be higher than the K_D_ of the simple BMP/BMPR interactions, indicating a lower affinity. In more details, the values of the two K_D_ values may be attributed to the values of two different types of binary interaction (BMP/ALK2 or BMP/ type-II BMPR), such as the interaction BMP-7/ALK2/BMPR-II where K_D_1 = 4.6 ± 0.7 nM and K_D_2 = 19.1 ± 6.4 nM (table.3), compared to 18.4 ± 3.8 nM for ALK2 and 5.5 ± 1.2 nM for BMPR-II in the binary experiments (Table.2). Nevertheless, this observation was not observed for other interactions. These results suggest the complexity of the interactions occurring on the surface.

**TABLE 3.**
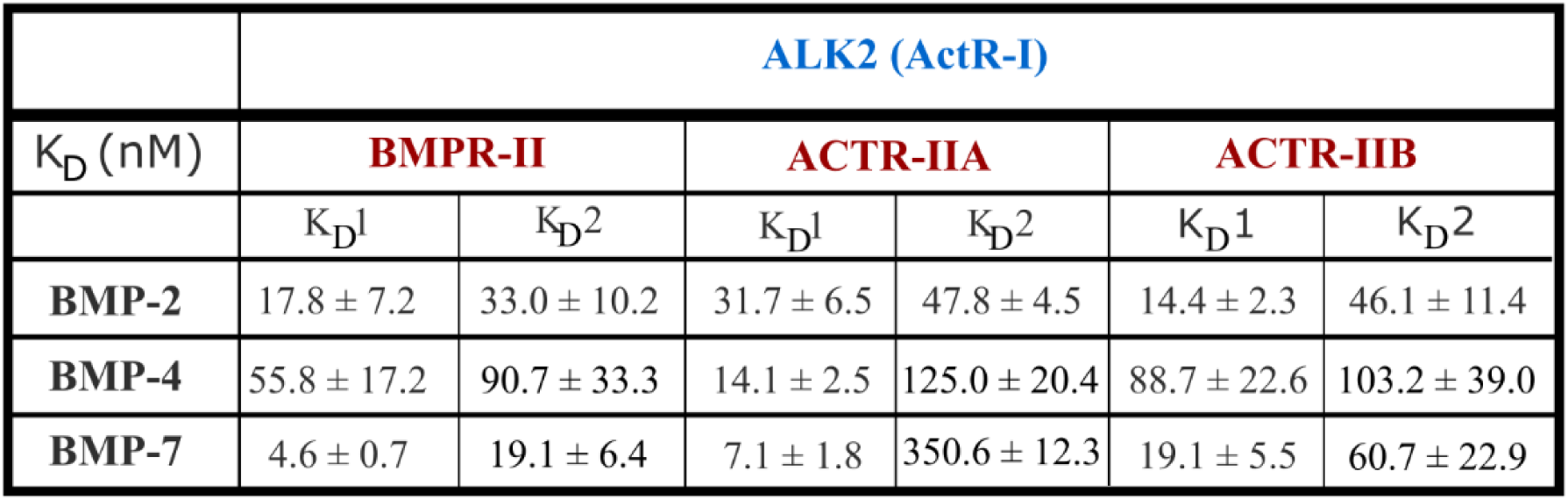
Binding affinities (K_D_) (nM) of BMP/BMPR-I/BMPR-II interactions. Table summarizing the K_D_1 and K_D_2 (nM) of the BMP/BMPR-I/BMPR-II interactions obtained from the kinetic experiments in a conformation where ALK2 and type-II BMPR are loaded sequentially. The error values represent the s.d (n=3).

|  | **ALK2 (ActR-I)** |  |  |  |  |  |
| --- | --- | --- | --- | --- | --- | --- |
| $K_D$ (nM) | **BMPR-II** | | **ACTR-IIA** | | **ACTR-IIB** | |
| | $K_{D1}$ | $K_{D2}$ | $K_{D1}$ | $K_{D2}$ | $K_{D1}$ | $K_{D2}$ |
| **BMP-2** | $17.8 \pm 7.2$ | $33.0 \pm 10.2$ | $31.7 \pm 6.5$ | $47.8 \pm 4.5$ | $14.4 \pm 2.3$ | $46.1 \pm 11.4$ |
| **BMP-4** | $55.8 \pm 17.2$ | $90.7 \pm 33.3$ | $14.1 \pm 2.5$ | $125.0 \pm 20.4$ | $88.7 \pm 22.6$ | $103.2 \pm 39.0$ |
| **BMP-7** | $4.6 \pm 0.7$ | $19.1 \pm 6.4$ | $7.1 \pm 1.8$ | $350.6 \pm 12.3$ | $19.1 \pm 5.5$ | $60.7 \pm 22.9$ |

## Discussion

In this study, we performed experiments using the dimeric form of BMPs and BMPRs as confirmed by gel electrophoresis migration analysis (Fig.SI.1). Our aim in using dimers of BMPs and BMPRs was to mimic the *in vivo* interactions since BMPRs were reported to be homodimer or heterodimers (type-I BMPR/type-II BMPR) at the cell surface, in contrast to TGFβR which appears to be homodimers in the absence of BMP-2 [51].

Using BLI, we quantified the binding affinities of the four BMPs with the eight different BMPRs in similar experimental conditions. As we showed with our SPR data, several BMPR/BMP couples (ALK3/BMP-2, ALK3/BMP-4 and ALK2/BMP-2) could not be analyzed by SPR using the same strategy as BLI with protein A coated sensors (Fig.SI.3), while BMP-9/ALK1 was detected (FIG.5). There may be non-specific adsorption of BMP-2 and 4 to protein A. The direct comparison of BLI versus SPR for the high affinity couple ALK1/BMP-9 showed that K_D_ measured by BLI was 15-fold lower than that measured by SPR (13.4 pM versus 200 pM) (Fig.5). These differences between both techniques may be explained by the physical and chemical differences of the techniques, since the thickness and composition of the sensor layers as well as the adsorption strategies are different. Another aspect to mention is the sensitivity of the method and the stability of the baseline signal, since the dissociation rate measured are sometimes at the limits of the instrument stability. Altogether, our experimental results show that the BLI technique is well adapted to gain quantitative information on a large range of BMP/BMPR couples.

To date, BMP-4 has barely been studied since it was often considered to exhibit a very close behavior to BMP-2 [5]. Our study first confirmed that both BMP-2 and BMP-4 bind to ALK3 and ALK6 with high affinity (Table.1), as already mentioned in the literature [6,30]. Additionally, our data reveals notable differences in the binding behaviors of BMP-2 and BMP-4. Indeed, BMP-2 binds to type-I BMPRs with a 3-fold higher affinity than BMP-4 (Table.2). Interestingly, the difference arises mainly from a difference in association rates to the receptors which was faster for BMP-2 than BMP-4, while the dissociation rates were similar (Fig.3). Notably, the strongest differences were observed for the type-II BMPR, with faster association rates for BMP-2 for all the three receptors, and faster dissociation rates for BMP-2 solely for BMRR-II and ACTR-IIA.

Our results are in agreement with previously published cellular data highlighting the distinct role of BMP-4 and BMP-2. One study examined their role in chondrocyte proliferation and found that the deletion of BMP-2 gene alone resulted in severe chondrodysplasia while the deletion of BMP-4 led to minor cartilage phenotype [52]. Likewise, a study using Xenopus embryos showed that the inhibition of BMP-4 or the use of a truncated ACTRII-A led to the similar phenotype for the formation of a secondary axis, while a defect in ACTRII-B caused an axial defect. As different phenotypes were observed, they concluded that BMP-4 does not transduce the biochemical signals via ACTR-IIB [53]. In acute myeloid leukemia, a distinct role of BMP-4 versus BMP-2 has been evidenced [54,55]: BMP-4 solely is involved as it activates a specific signaling pathway promoting immature resistant leukemic cells, which eventually leads to a relapse after treatment [54,55]. Furthermore, it has been demonstrated that BMP-4 mainly stimulated hematopoietic stem cell proliferation, differentiation and homing [56,57]. To note, these experimental results are not confirmed by structural data, although a study has shown that the crystal structures of the ternary complexes of BMP-2/ALK3/ACTR-IIA and BMP-2/ALK3/ACTR-IIB do not present significant structural differences [21]. In view of our findings regarding the specific differences between BMP-2 and BMP-4, it will be interesting to further evaluate their effects in different cell signaling contexts.

It is also noteworthy that the average binding of all BMPs (−2, 4, 7, 9) to ALK2 is in the same range ≈ 7-10 nM. In addition to the lower affinity of BMP-2 for type-II BMPRs compared to type-I BMPRs, we observed faster kinetic constants (k_a_, k_d_) for type-II BMPRs (Fig.3). This observation was previously reported and assumed to be the reason why BMP-2 and BMP-4 are recruited in a sequential order, with an initial binding to the higher affinity type-I BMPRs [21]. It may be interesting to further study BMP/ALK2 interactions in the context of the R206H mutation, which is associated to Fibrodysplasia Ossificans Progressiva (FOP): this mutation leads to the activation of BMP signaling in the absence of BMP and to an enhanced biochemical signal in the presence of BMP [58].

Our results showed that BMP-7 binds similarly to all ALKs with an affinity of ≈ 20 nM, in agreement with the literature review (Table.1), although the range of previously-reported K_D_ was large. With respect to type-II BMPRs, we found that BMP-7 binds with high affinities to the three type-II BMPRs, with a 5-fold higher affinity for ACTR-IIA (Table.2B). A previous study reported that BMP-7 signals through ACTR-IIA [59]. Notably, BMP-7 was also reported to induce chemotaxis in monocytic cells through BMPR-II and ACTR-IIA receptors, but not through ACTR-IIB [60].

Previous studies on BMP-9 have shown that it binds to ALK1 and ALK5 in endothelial cells [61–63], and to ALK1 and ALK2 in mesenchymal cells used for osteogenic differentiation [64]. Our data showing that BMP-9 binds ALK1 with a high affinity (0.2 ± 0.1 nM) and ALK2/ALK5 with a lower affinity (133.1 ± 35.1 nM and 51.0 ± 18.3 nM, respectively) (Table.2A) indicated that all these three ALK receptors are important in the signaling of BMP-9. The comparison of the structural data between both complexes BMP-2/ALK3-ECD/ACTR-2B-ECD and BMP-9/ALK1-ECD/ACTR-2B-ECD shows that ALK1 has a distinct interface with BMP-9, and presents several structural differences, compared to other type-I BMPRs. These structural disparities may well explain the low affinity of ALK1 for all the other BMPs [35], as seen in our data. Regarding the type-II BMPRs, former studies have shown that BMP-9 can bind to all of them [61,65]. Our results indicated that BMP-9 bound all type-II BMPR with a very high affinity (~ 0.8 to 1.7 nM) and with a slightly higher affinity for BMPR-II (0.8 ± 0.2 nM) (Table.2), in concert with the literature [35,36] (Table.SI.1).

Interestingly, our results showed that there is a binding of several BMPs to ALK5 (Fig 4 and Table.2A). We observed average affinities of BMP-2 (5.8 ± 1.1 nM), BMP-4 (21.9 ± 6.6 nM) and BMP-7 (22.6 ± 1.1 nM) to ALK5. Although ALK5 was considered to be mainly a TGFβR, our data show that several BMPs can bind to ALK5, which highlights its possible role in the BMP signaling pathway. Indeed, ALK5 was shown to inhibit BMP signaling mediated by ALK1 in the growth plate of cartilage [66]. It was also shown that different signaling through ALK1 and ALK5 regulate leptin expression in mesenchymal stem cells [16]. Last but not least, BMP-2 induces complex formation between ALK3 and ALK5 in cancer cells [45]. Further in vivo studies should aim to unravel a possible crosstalk between TGFβ/BMP pathways mediated by ALK5.

The use of a 2:1 heterogeneous ligand model to analyze the ternary complex interactions did not yield any improvement in the binding affinity compared to the bimolecular BMP/BMPR interactions, although such mechanism of cooperativity has been proposed. It was reported that BMP-7 affinity to ALK2 increases in the presence of ACTR-IIA [24,67]. Nonetheless, our data do not show any cooperativity between both types of BMPRs. This result agrees with the literature since a previous SPR study of BMP-7/ALK3/ACTR-IIA using a BMPR mix similarly reported limitation of the system in observing a cooperativity [28].

Our study highlighted the specific differences in BMP/BMPR interactions that could pave the way for future BMP signaling studies, with respect to BMP/TGFβ crosstalk and to the type of signaling pathway (SMAD versus non-SMAD) and to the specificities of the receptor (type I versus type II).

## Experimental procedures

### Protein and reagents

All used BMPs and extracellular domains (ECD) of the BMPR-FC chimeras were bought from Sigma Aldrich (Missouri, USA) and R&D systems (Minnesota, USA) respectively. BMP-2 and BMP-7 and BMP-9 are produced in Chinese hamster ovary (CHO) cells while BMP-4 was produced in *Escherichia coli*. The Anti-hIgG Fc (AHC) capture biosensors were purchased from ForteBio (California, USA) and the protein A coated chips were purchased from GE Healthcare Life Sciences. The buffer was made of 20 mM HEPES at pH 7.4 with 150 mM NaCl (name hereafter Hepes-NaCl) and 0.02% tween while the regeneration buffer was made of 10 mM glycine at pH 1.7 (named hereafter regeneration buffer). They were all prepared in-house.

### Kinetics interaction experiments

All the BLI experiments were performed using an OctetRED96e apparatus from Pall/FortéBio (California, U.S) and data were recorded with the manufacturer software (Data Acquisition v11.11). All proteins were solubilized following the supplier instruction in Hepes-NaCl buffer. The analysis protocol was adapted from previous studies [30,35]. In details, prior any capture, the BMPR-Fc samples were first diluted in the Hepes-NaCl buffer. For the association phase, the BMPs were diluted in 2-fold serial dilutions in Hepes-NaCl buffer. 0.2 ml of each sample and buffer were disposed in wells of black 96-well plates (Nunc F96 MicroWell, Thermo Fisher Scientific), maintained at 25°C and agitated at 1000 r.p.m. the whole time. Prior each assay, all biosensors were pre-wetted in 0.2 ml of Hepes-buffer for 10 min, followed by monitored equilibration for 60 or 120 s. Anti-hIgG Fc (AHC) capture biosensors (FortéBio) were loaded with each ligand for 200 s until to reach a spectrum shift between 0.8 and 1.1 nm depending of BMPR-Fc, followed by an additional equilibration step of 60 s or 120 s in Hepes-NaCl buffer. Association phases were monitored during dipping the functionalized biosensors in analyte solutions of different concentrations between 2 and 80 nM for 400 s, and the dissociation phases in the buffer for 400 s. To assess and monitor analyte unspecific binding, blank biosensors were treated with the same procedures but replacing the ligand solutions by analysis buffer. All sensors were fully regenerated between experiments with different BMPRs by dipping for 30s in regeneration buffer. All measurements were performed three times in independent experiments.

Kinetics data were analyzed using the manufacturer software (Data analysis HT v11.1). The “blank” signal from the biosensor in the presence of the Hepes-NaCl buffer was subtracted from the signal obtained from each functionalized biosensor and each analyte concentration. The kinetic signals were then fitted using a global/local method and 1:1 Langmuir or 2:1 heterogeneous ligand model. Affinity constants were calculated from the ratio k_d_/k_a_ values. The reported values are given as mean ± SD obtained from three independent experiments.

### Surface plasmon resonance experiments

All surface plasmon resonance experiments were performed using a Biacore T200 apparatus (GE Healthcare Life Sciences/Biacore, Illinois, U.S) and data were recorded using the manufacturer software (Biacore control software v2.0). All protein samples were solubilized following the supplier instruction in analysis buffer prior any experiment. Prior to capture, the BMPR-Fc samples were first diluted in analysis buffer. For association phase, BMP samples were diluted at concentration between 0.2 and 6.4 nM in 2-fold serial dilutions in the Hepes-NaCl buffer. Sensor chips and system were pre-equilibrated in Hepes-NaCl buffer prior any injection. The protein A sensor chips (GE Healthcare Lifesciences) were loaded by injecting each ligand for 100 s until to reach a signal level between 100 and 120 arbitrary response units (R.U.) depending of BMPR-Fc, followed by an additional equilibration step of several minutes in analysis buffer. Association phases were monitored during injections over the functionalized surfaces of analyte solutions of different concentrations for 300 s, and the dissociation phases of analysis buffer for 300 s. To assess and monitor analyte unspecific binding, blank surfaces were treated with the same procedures but replacing the ligand solutions by analysis buffer. All surfaces were fully regenerated between experiments with different BMPR-Fc by injecting for 30s regeneration buffer. Two independent experiments were performed. Kinetic data were processed with the manufacturer software (Biacore Evaluation software v3.1). Signals from the reference surface were subtracted from the signals obtained from each functionalized ship. Resulting specific kinetics signals were then fitted using the 1:1 Langmuir model. Affinity constants were calculated from the ratio K_D_/k_a_ values. Reported values are obtained by averaging the values obtained from the replicates and reported errors as the standard deviation.

## Supporting information

Khodr-supplemental file

## Author contributions

C.P conceived the idea. C.P, P.M, VK and E.M designed the experiments. V.K and P.M performed the experiments and collected the data. VK, P.M, C.P and J.B.R analyzed the data. C.P acquired the funding. V.K., P.M, J.B.R and CP wrote the manuscript. E.M. reviewed and edited the manuscript.

## Acknowledgments

The authors acknowledge Anne Chouquet from the ISBG platform for her help and Marianne Weidenhaupt (INPG) for discussions regarding the immobilization protocol. We are grateful to Sabine Bailly and Corinne Albiges-Rizo for fruitful discussions and advices.

## Funding and additional information

The study was supported by Agence Nationale de la Recherche (ANR CODECIDE, ANR-17-CE13-022) to C.P and by the Fondation Recherche Medicale (FRM, grant DEQ20170336746) to C.P. VK was supported by a PhD fellowship from Grenoble Institute of Technology. This work used the platforms from Grenoble Instruct-ERIC Center (ISBG: UMS 3518 CNRS-CEA-UGA-EMBL), within the Grenoble Partnership for Structural Biology (PSB), supported by FRISBI (ANR-10-INBS-05-02) and GRAL, financed within the University Grenoble Alpes graduate school (Ecoles Universitaires de Recherche) CBH-EUR-GS (ANR-17-EURE-0003).

## Supporting information

**SI TABLE 1**. Detailed literature study table summarizing the K_D_ (nM) of the BMP/type-I and type-II BMPR interaction couples.

**SI FIGURE 1.** Image of a gel electrophoresis showing all the used BMPs, ALKs and type-II BMPR.

**SI TABLE 2**. Detailed kinetic tables indicating the K_D_ (nM), k_a_ (M^−1^.s^−1^), k_d_ (s^−1^) and R^2^ of the BMP/type-I interaction couples.

**SI TABLE 3**. Detailed kinetic tables indicating the K_D_ (nM), k_a_ (M^−1^.s^−1^), k_d_ (s^−1^) and R^2^ of the BMP/type-II interaction couples.

**SI FIGURE 2.** Binding kinetics between non-glycosylated and glycosylated BMP-4 with ALK3.

**SI FIGURE 3.** SPR binding curve for BMP-2/ALK3.

**SI FIGURE 4.** BLI binding kinetics of BMP-2 with ALK2 and ACTR-IIB in two confirmations using 1:1 fit.

