## Supplementary material for "High throughput measurements of BMP/BMP receptors interactions using bio-layer interferometry": Khodr-supplemental file

**SI TABLE 1. Detailed literature study table summarizing the $K_D$ (nM) of the BMP/type-I and type-II BMPR interaction couples**. The experiments are usually performed in two different configurations: in red, when the BMP-R is immobilized; in blue when the BMP is immobilized, using SPR and BLI as techniques and different loading strategies.

| $K_D$ | Immo. | Tech. | ALK1 | ALK2 (ActR-I) | ALK3 (BMPR-IA) | ALK5 (TGFβR-I) | ALK6 (BMPR-IB) | BMPR-II | ACTR-IIA | ACTR-IIB | Adsorption tech. | Ref. |
|---|---|---|---|---|---|---|---|---|---|---|---|---|
| BMP-2 | BMPR | SPR | | | 1 nM | | 11 nM | 100 nM | 50 nM | | Biotin/ Streptavidin | Kirsch,T. et al (2000) |
| | | SPR | | | | | | | 38 nM | | Amine coupling | Greenwald, J et al (2003) |
| | | SPR | | | | | | | 38 - 80 nM | | Amine coupling | Allendorph, G. et al (2006) |
| | | SPR | | | 2.6 nM | | | | 47.5 nM | 36.1 nM | Amine coupling | Allendorph, G. et al (2007) |
| | | SPR | | | 0.7 nM | | 2.7 nM | | | | Biotin/ Streptavidin | Kotzsch, A et al (2008) |
| | | SPR | | | 0.8 nM | | 2.7 nM | 45±20 nM | 14 nM | 6.3 nM | Biotin/ Streptavidin | Heinecke, K. et al (2009) |
| | | SPR | | | 3.75 pM | | 2.48 nM | | | | BMPR-Fc on anti-Fc | Yamawaki, K. et al (2016) |
| | | BLI | | | 1.1 nM | | 1.1 nM | 26.7 nM | 52.7 nM | 8 nM | BMPR-Fc on anti-Fc | Seeherman, H. et al (2019) |
| | BMP | SPR | | | | | | | 5.4 µM | | Amine coupling | Greenwald, J et al (2003) |
| | | SPR | | >400 µM | 10 µM | | 95 µM | | 3.8 µM | | Amine coupling | Saremba, S. et al ( 2007) |
| | | SPR | | | 48 nM | | 350 nM | | | | Biotin/ Streptavidin | Heinecke, K et al (2009) |
| | | SPR | | | 330 nM | | | | | | Amine coupling | Mahlawat, P. et al (2012) |
| BMP-4 | BMPR | SPR | | | 56.4 pM | | 221 pM | | | | See above | Yamawaki, K. et al (2016) |
| | BMP | SPR | | | 47 nM | | | | | | Amine coupling | Hatta, T. et al (2000) |
| | | SPR | | | 1.2 - 9.6 nM | | | | | | Amine coupling | Szlama, G et al (2010) |
| BMP-7 | BMPR | SPR | | | | | | | 1.2 nM | | See above | Greenwald, J et al (2003) |
| | | SPR | | | | | | | 1 nM | | See above | Allendorph, G. et al (2006) |
| | | SPR | | | 1680 nM | | | | 3.49 nM | 7.72 nM | See above | Allendorph, G. et al (2007) |
| | | SPR | | > 500 nM | 58 nM | | 9 nM | 25 nM | 5.1 nM | 6.5 nM | See above | Heinecke, K. et al (2009) |
| | | SPR | | | 1.95 nM | | 268 pM | | | | See above | Yamawaki, K. et al (2016) |
| | BMP | SPR | | 143 µM | | | | | 1.7 µM | | See above | Greenwald, J et al (2003) |
| | | SPR | | 55 µM | 10 µM | | 1 µM | | 0.9 µM | | See above | Saremba, S. et al ( 2007) |
| | | SPR | | | 1900 nM | | 750 nM | | | | See above | Heinecke, K et al (2009) |
| BMP-9 | BMPR | SPR | 45 pM | | | | | 630 pM | 6.43 nM | 21.7 pM | BMPR-Fc on anti-Fc | Townson, S. et al (2012) |
| | | SPR | 20 - 45 nM (monomer)/ 2-3 nM (dimer) | | | | | | | | Amine coupling | Mahlawat, P. et al (2012) |
| | | SPR | <8 pM | | | | | 3.3 nM | 42.7 nM | 1.4 nM | BMPR-Fc on anti-Fc | Kienast, Y. (2016) |
| | | SPR | 71.6 pM (monomer) / 48.1 pM (dimer) | | | | | | | | Amine coupling | Salmon, R. (2020) |

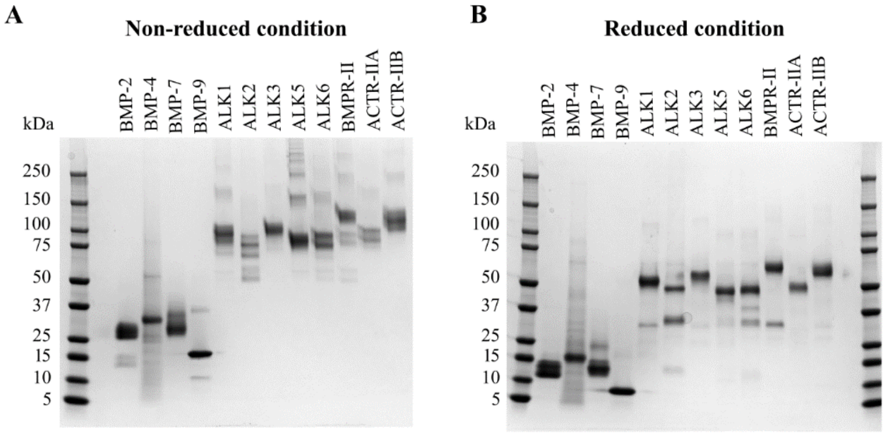


**SI FIGURE 1. Image of a gel electrophoresis showing all the used BMPs, ALKs and type-II BMPR.** The proteins were tested in **A)** non-reducing and **B)** reducing conditions.

**SI TABLE 2. Detailed kinetic tables indicating the $K_D$ (nM), $k_a$ ($M^{-1}.s^{-1}$), $k_d$ ($s^{-1}$) and $R^2$ of the BMP/type-I BMPR interactions.** The error values represent s.d (n=3).

| ALK / BMP | ALK1 | | | | ALK2 (ACTR-I) | | | |
|---|---|---|---|---|---|---|---|---|
| | kinetic KD | ka (1/M.s) ($10^5$) | kdis (1/s) ($10^{-3}$) | $R^2$ | kinetic KD | ka (1/M.s) ($10^5$) | kdis (1/s) ($10^{-3}$) | $R^2$ |
| BMP-2 | 13,0 ± 1.6 | 2.5 ± 1.3 | 3.2 ± 1.2 | 0.94 ± 0.02 | 7.0 ± 2.3 | 7.4 ± 2.3 | 5,0 ± 1.3 | 0.93 ± 0.01 |
| BMP-4 | 55.4 ± 4.0 | 0.74 ± 0.24 | 4.1 ± 1.5 | 0.95 ± 0.002 | 10.5 ± 3.8 | 2.3 ± 1.7 | 2.6 ± 1.6 | 0.92 ± 0.04 |
| BMP-7 | 23.1 ± 2.1 | 1.6 ± 0.2 | 3.7 ± 0.4 | 0.95 ± 0.02 | 18.4 ± 2.4 | 2.0 ± 0.5 | 3.6 ± 0.2 | 0.96 ± 0.04 |
| BMP-9 | 0.17 ± 0.08 | 15,0 ± 3.5 | 0.2 ± 0.1 | 0.99 ± 0.01 | 133.1 ± 35.1 | 1.2 ± 0.9 | 16 ± 3 | 0.93 ± 0.02 |

| ALK / BMP | ALK3 (BMP-R-IA) | | | | ALK5 (TGFβR-I) | | | |
|---|---|---|---|---|---|---|---|---|
| BMP-2 | 0.21 ± 0.03 | 11.3± 0.6 | 0.26 ± 0.05 | 0.99 ± 0.003 | 5.8 ± 1.1 | 4.2 ± 0.4 | 2.4 ± 0.5 | 0.95 ± 0.03 |
| BMP-4 | 1.7 ± 0.5 | 7.3 ± 1.6 | 1.5 ± 0.2 | 0.97 ± 0.03 | 21.9 ± 6.6 | 2.05 ± 0.7 | 4.3 ± 1.0 | 0.94 ± 0.02 |
| BMP-7 | 19.0 ± 2.1 | 1.2 ± 0.4 | 2.3 ± 0.4 | 0.96 ± 0.02 | 22.6 ± 1.1 | 1.6 ± 0.5 | 3.5 ± 0.3 | 0.96 ± 0.03 |
| BMP-9 | Very low signal | Very low signal | Very low signal | Very low signal | 51 ± 18.3 | 0.6 ± 0.06 | 3.0 ± 0.8 | 0.94 ± 0.004 |

| ALK / BMP | ALK6 (BMPR-IB) | | | |
|---|---|---|---|---|
| BMP-2 | 0.5 ± 0.1 | 19.1 ± 3.5 | 0.9 ± 0.1 | 0.96 ± 0.02 |
| BMP-4 | 3.0 ± 0.3 | 5.1 ± 0.5 | 1.6 ± 0.3 | 0.95 ± 0.04 |
| BMP-7 | 14.5 ± 1.2 | 1.8 ± 0.1 | 2.7 ± 0.1 | 0.98 ± 0.005 |
| BMP-9 | Very low signal | Very low signal | Very low signal | Very low signal |

**SI TABLE 3. Detailed kinetic tables indicating the $K_D$ (nM), $k_a$ ($M^{-1}.s^{-1}$), $k_d$ ($s^{-1}$) and $R^2$ of the BMP/type-II BMPR interactions.** The error values represent s.d (n=3).

| ALK / BMP | BMPR-II | | | | ACTR-IIA | | | |
|---|---|---|---|---|---|---|---|---|
| | kinetic KD | ka (1/M.s) ($10^5$) | kdis (1/s) ($10^{-3}$) | $R^2$ | kinetic KD | ka (1/M.s) ($10^5$) | kdis (1/s) ($10^{-3}$) | $R^2$ |
| BMP-2 | 5.4 ± 0.8 | 12 ± 5.3 | 6.4 ± 2.3 | 0.96 ± 0.03 | 6.08 ± 1.2 | 13.5 ± 5.8 | 8.1 ± 1.6 | 0.95 ± 0.01 |
| BMP-4 | 56,0 ± 6,0 | 0.9 ± 0.2 | 4.8 ± 0.4 | 0.96 ± 0.01 | 21.4 ± 3.7 | 1.5 ± 0.5 | 3.2 ± 0.6 | 0.97 ± 0.01 |
| BMP-7 | 5.5 ± 1.2 | 4.3 ± 0.4 | 2.3 ± 0.5 | 0.99 ± 0.01 | 1.3 ± 0.3 | 9.9 ± 2.4 | 1.3 ± 0.2 | 0.98 ± 0.01 |
| BMP-9 | 0.8 ± 0.2 | 13.6 ± 4.0 | 1.2 ± 0.1 | 0.99 ± 0.43 | 1.7 ± 0.1 | 35.8 ± 9.1 | 6.0 ± 0.7 | 0.93 ± 0.03 |

| ALK / BMP | ACTR-IIB | | | |
|---|---|---|---|---|
| BMP-2 | 6.3 ± 3.4 | 8.5 ± 3.1 | 5.3 ± 0.6 | 0.94 ± 0.01 |
| BMP-4 | 26.0 ± 0.5 | 2.4 ± 0.6 | 6.1 ± 1.5 | 0.95 ± 0.02 |
| BMP-7 | 7.1 ± 0.7 | 4.0 ± 0.3 | 2.8 ± 0.4 | 0.99 ± 0.01 |
| BMP-9 | 1.4 ± 0.4 | 22.4 ± 4.6 | 2.6 ± 0,3 | 0.95 ± 0.04 |

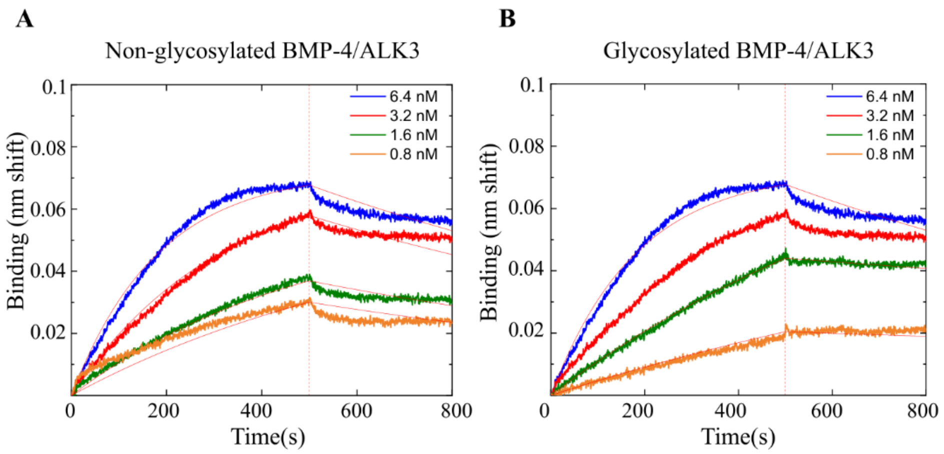


**SI FIGURE 2. Binding kinetics between non-glycosylated and glycosylated BMP-4 with ALK3**. **A)** non-glycosylated BMP-4 and **B)** glycosylated BMP-4.

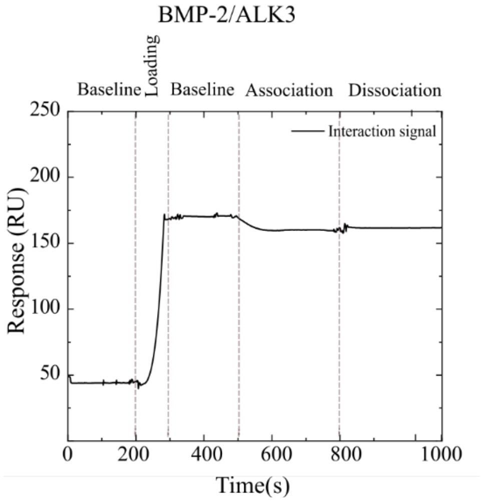


**SI FIGURE 3. SPR binding curve for BMP-2/ALK3**. The association signal was undetectable and even slightly decreased after background subtraction, showing that the interaction was not detectable.

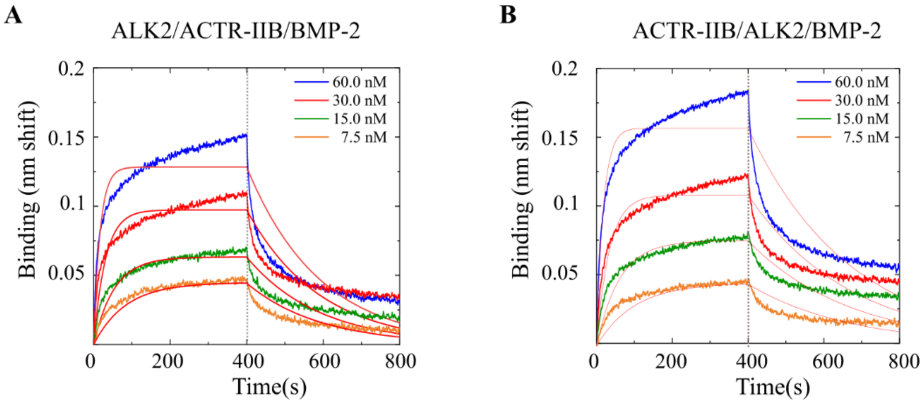


**SI FIGURE 4. BLI binding kinetics of BMP-2 with ALK2 and ACTR-IIB in two confirmations using 1:1 fit. A)** ALK2/ACTR-IIB/BMP-2 with ALK2 loaded first and **B)** ACTR-IIB/ALK2/BMP-2with ACTR-IIB loaded first. The data were analyzed using a 1:1 fit.
